# Zinc restriction promotes β-cell hyper-hormonemia and endocrine pancreas degeneration in mice

**DOI:** 10.1101/616201

**Authors:** Tháyna Sisnande, Cleverton K. Lima, Dayana Cabral da Silva, Thayana Moulin Beninatto, Natália Leão Alves, Mariana J. Amaral, Leandro Miranda-Alves, Luís Maurício T. R. Lima

## Abstract

Zinc is a key component of proteins, including interaction with varying pancreatic hormones, including insulin and amylin. Zinc is key in insulin crystallinity in ZnT8 knock-out mice models, although the dietary role of zinc restriction over both energetic metabolism and ß-pancreatic hormonemia and morphology remained unexplored. We aimed to test whether dietary zinc restriction on swiss male mice would impact over endocrine pancreas and metabolic phenotype. We evaluated the role of dietary zinc restriction on ß-pancreatic hormonemia on non-transgenic Swiss male mice weaned onto a control or low-zinc diet for 4 weeks. Growth, glycemia, insulinemia, amylinemia and pancreatic islet were smaller in intervention group despite insulin crystallinity in secretory granules. We have found overlabelling for insulin, amylin and toxic oligomers in apoptotic pancreatic islet. High production of β-pancreatic hormones in zinc-restricted animals counteract the decreasing islet size due to their apoptotic cells. We conclude that zinc deficiency is sufficient to promote islet β-cell hormonal disruption and degeneration.

## INTRODUCTION

Diabetes is a cluster of endocrine and metabolic disorders which have in common the inability to properly produce, secrete and/or respond to the β-pancreatic hormones insulin and amylin [1]. Type 1 diabetes (T1D) is characterized by loss of pancreatic β-cells mass and typically accompanied by the generation of autoantibodies, leading to the deficiency of both insulin and amylin [2]. Type 2 diabetes is due to insulin resistance, which results in normoglycemic hyperinsulinemia and hyperamylinemia at subclinical phases, prior to the hyperglycemic phase which accompanies β-cells destruction [3].

Both Type 2 diabetes (T2D) and T1D have a significant genetic component [4]. However, the increasing global trends and prevalence of T2D and other non-communicable diseases is not likely to be supported by the genetic background solely. Instead, on top of an increased genetic predisposition for T2D, several components may exert key roles in the ontogeny of the disease, including environmental factors, endocrine chemical disruptors, lifestyle and/or micronutrient imbalance [5,6].

Changes on lifestyle, which includes a higher consumption of micronutrient-poor food, has been associated to the increased incidence of non-communicable disorders [7]. Evidences suggest that essential micronutrients, in particular zinc, are important players in the onset and/or progression of different amyloid diseases [8–10]. Indeed, zinc supplementation has been shown to improve the clinical symptoms of patients [11–13]. Zinc participates on manifold biological processes, such as DNA replication and protein synthesis [14], changes on secretion and action of growth hormone [15], outcome on insulin-like growth factor-1 [16], protein stabilization [17] and others.

Zinc is the second most abundant micronutrient on organism, being found in about 10% of human proteins [18]. Zinc-related disorders can be either by dietary deficiency (ICD-10-CM Diagnosis Code E60) or metabolism disorders (ICD-10-CM Diagnosis Code E83.2). The diary dose recommended in adults is about 10 mg [19], but the total intake, bioavailability and bioaccessibility of this metal can be disturbed by food pattern and lifestyle [20]. Independent of the economic power, occidental societies have increased the consumption of micronutrient-poor food, restricted in zinc (NHANES III). Such pattern has been associated to increased propensity for metabolic, degenerative and neurodegenerative diseases [21–24]. Disruption of cell proteostasis can be mediated by amyloid proteins, and in this context oligomerization and aggregation/fibrillation would be secondary to micronutrients imbalance [25–27]. Nevertheless, zinc supplementation has been shown to bring clinical benefits to diabetic patients [28], which rise the question on the role of metal on islet amyloidogenesis and proteostasis.

In pancreatic β-cells, zinc is found at millimolar levels in secretory granules, and associated to crystalline insulin [29]. Zinc transporter ZnT8-knockout (ZnT8-/-) mice loss insulin crystallinity, and under a high-fat diet the ZnT8(-/-) mice became glucose intolerant or diabetic [30].

Amylin (also known as islet amyloid polypeptide, IAPP) plays role in metabolism and glucose homeostasis [31,32]. In addition, amylin is co-produced and co-secreted with insulin by β-cells [33]. The deficiency of insulin on type 1 diabetes *mellitus* (T1DM) and advanced T2D leads to consequent reduction of circulating amylin, while plasma levels are elevated in insulin-resistant conditions such as obesity and impaired glucose tolerance [34,35]. It is known that human amylin forms amyloid fibrils [36,37], but the aggregation of murine variant still under discussion. Our group demonstrated that murine amylin and amylin analogue pramlintide can aggregate *in vitro* [38–40]. Toxic oligomers in pancreatic islets can be found in non-transgenic mice [41]. Similarly, Leyva-García and colleagues found responsive material to antibody anti oligomer-like immunoreactive material (“anti-amyloid oligomer”, A11 antibody) on streptozotocin-induced mice [42]. These findings corroborate with previous identification of aggregated amylin on diabetic and elder nondiabetic individuals [43].

Zinc has been described as a partner in amylin oligomerization and fibrillation, both human [44–47] and murine [38] The zinc content on β-pancreatic granule has been shown to be reduced in diabetic patients (Sjögren *et al.* 1988) compared to healthy individuals [49], which rise the hypothesis of the role of pancreatic zinc in triggering the instability of amylin *in vivo*.

Motivated by animal, clinical and also epidemiological data showing improvements of diabetic panel by zinc supplementation [28], we decided to investigate the implications of reduction in dietary zinc on non-transgenic mice metabolism and endocrine pancreatic function.

## MATERIAL AND METHODS

### Material

Rodent purified diets (***Supplemental Material#1 to #3***) were formulated based on the AIN93G [50], with casein replaced by hen egg white solids (named “Albumina”, by Salto’s LTDA or Netto Alimentos LTDA; ***Supplemental Material#4***) as recommended by [50]. Control diet was formulated with the AIN93G mineral mix (AIN93G-MX-control) and the zinc-deficient diet was formulated with a mineral mix without zinc carbonate (AIN93mG-MX-ZnDef). Diets batches were prepared both by Rhoster Ind. Com. LTDA (São Paulo) and by one of our labs (*pbiotech*; with diet components from Farmos Com. Ind. LTDA, Rio de Janeiro). The Zn content in the diets were determined by mineral analysis with both a contract research organization CRO (Laboratório de Ciência e Tecnologia, São Paulo, Brazil; ***Supplemental Material #5 and #6***) and Empresa Brasileira de Produtos Agropecuários (EMBRAPA, Rio de Janeiro; not shown). All reagents were from analytical grade. All reactants were used as received.

### Animal experimentation

This study was approved by the Institutional Bioethics Committee of Animal Care and Experimentation at the UFRJ (CCS, UFRJ, 011/2015). Swiss male mice were randomly divided and fed with iso or hypozincemic diet for 4 weeks (n=15 per group). Animals were housed in a temperature-controlled room with a light-dark cycle of 12 h.

Once a week, at about same day time (1 pm) mice were evaluated for body mass and non-fasting blood glycemia. The glycemia was measured from the tail tips of conscious and unrestrained mice using pre-calibrated point-of-care glucometers (Accu-Chek Active, Roche Diagnostics). At 4^th^ week, ketosis (β-hydroxy-butyrate, BHB) was evaluated (Precision Xtra Blood Glucose and Ketones Bodies monitoring system, Abbot, USA) and fed animals were killed, and glycated haemoglobin was measured using A1CNow^®^+ Multi-Test A1c System (PTS Diagnostics, Indianapolis, IN, USA). Subsequently to cardiac puncture, samples were rapidly homogenised with EDTA by inversion 5 times, centrifuged at 2000 × g for 10 min at room temperature and the plasma was separated and stored at −20°C until analysis on clinical chemistry analyzer (Beckman Coulter AU5800, Beckman Coulter Inc. Brea, CA; HUCFF-UFRJ), using kinetic colorimetric assays for α-amylase (Beckman OSR6006) and lipase (Beckman OSR6230). Statistical analysis was performed using one-way analysis of variance (ANOVA) with Bonferroni test as post hoc analysis by using GraphPad Prism ver 5.01 (GraphPad Software). A *p*-value of <0.05 was considered significant.

### Insulin and amylin measurement assay

Insulin and amylin concentration in plasma were determinate by ELISA (Insulin, Cat # EZRMI-13K and Amylin, Cat # EZHA-52K, Merck Millipore) following manufacturer manual. Statistical analysis was performed using unpaired t-test with as post hoc analysis by using GraphPad Prism ver 5.01 (GraphPad Software). A p-value of <0.05 was considered significant.

### Histological analysis

Pancreas was isolated, washed on cold PBS buffer (137 mM NaCl, 2.7 mM KCl, 8 mM Na_2_HPO_4_ and 2 mM KH_2_PO_4_, pH 7.4) and dived into three equal pieces to three different fixation mediums. Pancreas was fixed on formalin 10% buffer pH 7.4, and tissues were processed and embedded on paraffin. Slices of 3 µm were obtained and analyzed by optical microscopy (Olympus microscope, model BX60F5, Olympus optical, Co. Japan. Coupled CCD digital camera QImaging Retiga 2000R High-Sensitivity IEEE 1394 FireWire. San Diego, CA. USA). Hematoxylin-eosin (HE) stained sections were used to measure Langerhans islets area using ImageJ [51]. Picrosirius red staining were used to quantify total collagen on pancreatic tissue using Image Pro Plus (Media Cybernetics Inc. version 6.00.260). Sirius red solution were prepared by dissolving 500 mg of Sirius Rred (Direct Red 80; Cat # 365548, Sigma Aldrich) with 500 mL saturated solution of picric acid. Paraffin sections were dewaxed in xylol, hydrated in graded ethanol, distilled water and stained with hematoxylin. Sections were incubated on picrosirius red staining for 1h and washed twice in acidic water (acetic acid 0.005% in water). Statistical analysis was performed using unpaired t-test using GraphPad Prism version 5.01 (GraphPad Software). A p-value of <0.05 was considered statistically significant.

### Immunoassay

Pancreatic tissue was collected and fixed in paraformaldehyde 4%. Paraffin sections were dewaxed in xylol, hydrated in graded ethanol and distilled water. To antigen retrieval were used 10 mM citrate buffer, pH 6.0, 35 min at 95°C. Thereafter, the slices were rinsed in PBS pH 7.4 or 10 mM Tris pH 7.4 and the endogenous peroxidase was blocked with 3% hydrogen peroxide in methanol for 20 min. After blocking with PBS solution containing 5% bovine serum albumin, 0.25% Triton X-100, 0.025% tween and 0.1% gelatin. Sections were incubated overnight in a humid chamber (4°C) with primary anti-insulin antibody (Insulin Antibody #4590 4590S, Cell Signaling), anti-oligomer antibody (anti prefibrillar oligomer-like material; Cat #AB9234, Merck-Millipore) or anti-murine amylin (in-house derived, from rabbits, produced by EJAPC) at 1:100 dilution for peroxidase assay. After washing with 0. PBS pH 7.4, plus 0.25% Tween X-100, tissues were incubated for 1 hour at room temperature with Histofine^®^ Simple Stain MAX PO (code 414341F, Nichirei Biosciences Inc. Tokyo. Japan) and counterstained with Harris’s hematoxylin. Additional images of representative histological data can be found in **Supplemental Material**.

### Transmission electron microscopy (TEM)

For TEM, pancreatic tissues were cut into small pieces (about 3 mm x 3 mm), fixed during 4 days in Karnowskýs buffer (made fresh, 2.5% glutaraldehyde, 4% paraformaldehyde, sodium cacodylate 0.1 M, pH 7.2) at room temperature. Tissues were washed in 0.1 M sodium cacodylate pH 7.2, post fixed for 40 min in 1% OsO_4_ in cacodylate buffer containing 1.25% K₃[Fe(CN)₆] and 5 mM CaCl_2_ and dehydrated in an ascending acetone series. Subsequently, tissues were embedded in Epoxi resin (Cat# 14120, Electron Microscopy Science, Hatfield, PA). Semithin section (2.5 µm) were obtained with glass knife and 1% toluidine blue solution staining for islet localization. Finally, thin sections (60-70 nm) were contrasted with uranyl acetate, lead citrate. Images were acquired in a Zeiss 900 transmission electron microscopy (Carl-Zeiss, Oberkochen, Germany).

### TUNEL assay

The detection of apoptotic cells was based on the nick-end (TUNEL) labelling method using the ApopTag® *in situ* detection kit (S7100, Merck Millipore, USA), according to the manufacturing protocol.

## RESULTS

### Experimental model of zinc dietary restriction in non-transgenic mice

Weaning Swiss male mice were submitted to either a control diet or a diet restricted in the total amount of zinc (38 mg/kg intervention vs 11 mg/kg control, respectively). Mice were followed for about 4 weeks. The time-course of body mass gain was less prominent in the intervention group (**Fig. 1A**). The animals submitted to zinc restriction also showed smaller body length, reduced in about 20 % compared to control, as indicated by the shorter tibia length (**Fig. 1B**). The manifestation of restricted growth and body mass due to restriction in dietary zinc are in good agreement with previous observation from experimentation in rats [52–54], confirming the strength of the present intervention.

**Fig 1.**
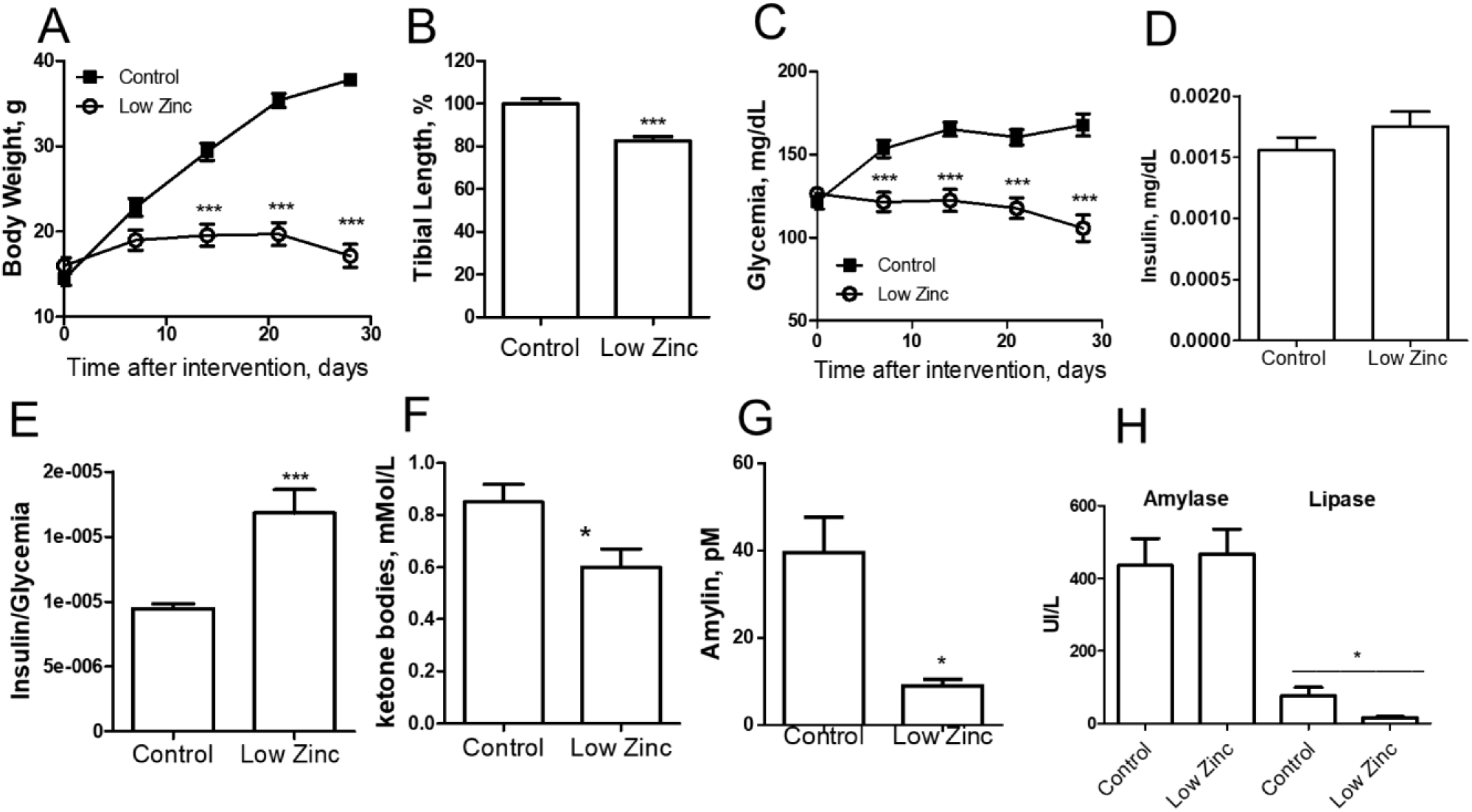
Effects of moderate zinc restriction in non-transgenic mice. Weanling Swiss male mice were submitted on a moderate zinc diet for 28 days, animals exhibit restriction in weight gain (A) and growth (B). Lower glucose levels were observed in low zinc diet animals in comparison to control group (C). Insulin plasmatic levels had no difference between groups (D), although zinc restriction diet presented insulin resistance (E). Ketone bodies were reduced in intervention group (F) and circulating amylin is decreased on the second group (G). N = 15. * *p*< 0.05, *** *p*< 0.0001.

During growth we observed a normal progression of the glycemia in control group, which was halted in the intervention group (**Fig. 1C**). Fasting insulinemia showed no difference between groups after 4 weeks dietary intervention (**Fig. 1D**). Despite the lower glycemia in the intervention group, they showed a higher insulin/glycemia ratio compared to control group (**Fig. 1E**), suggestive of insulin resistance. Animals submitted to zinc-restricted diet showed higher circulating β-hydroxy-butyrate (BHB) levels (**Fig. 1F**), lower glycated hemoglobin (A1C) (**Fig. S1**) and lower circulating amylin (**Fig. 1G**) compared to control group. No major alteration in exocrine pancreas took place, as judged by equivalent pancreatic amylase and only minor reduction of pancreatic lipase (**Fig. 1H**). Although no significant difference in plasmatic zinc have been observed between groups (**Fig S2**), these collective data indicate a major effect of zinc restriction over glycemic control and pancreatic endocrine function.

### Zinc restriction induces endocrine pancreas disturbance

Pancreatic histological analyses of both groups (**Fig. 2**) revealed an infiltration of adipose tissue on intervention group (arrows). Another pronounced difference between the groups was the architecture of islets of intervention group, as depicted by the non-typical form of islets with undefined edges (**Fig. 2B**). Additionally, the pancreatic islet area was also smaller in intervention group, about one third the size of control group (**Fig. 2C**), disproportional to the reduced body size (about 80 % of control group – **Fig. 1B**), indicating that other underlying mechanisms are behind the reduced islet size.

**Fig 2.**
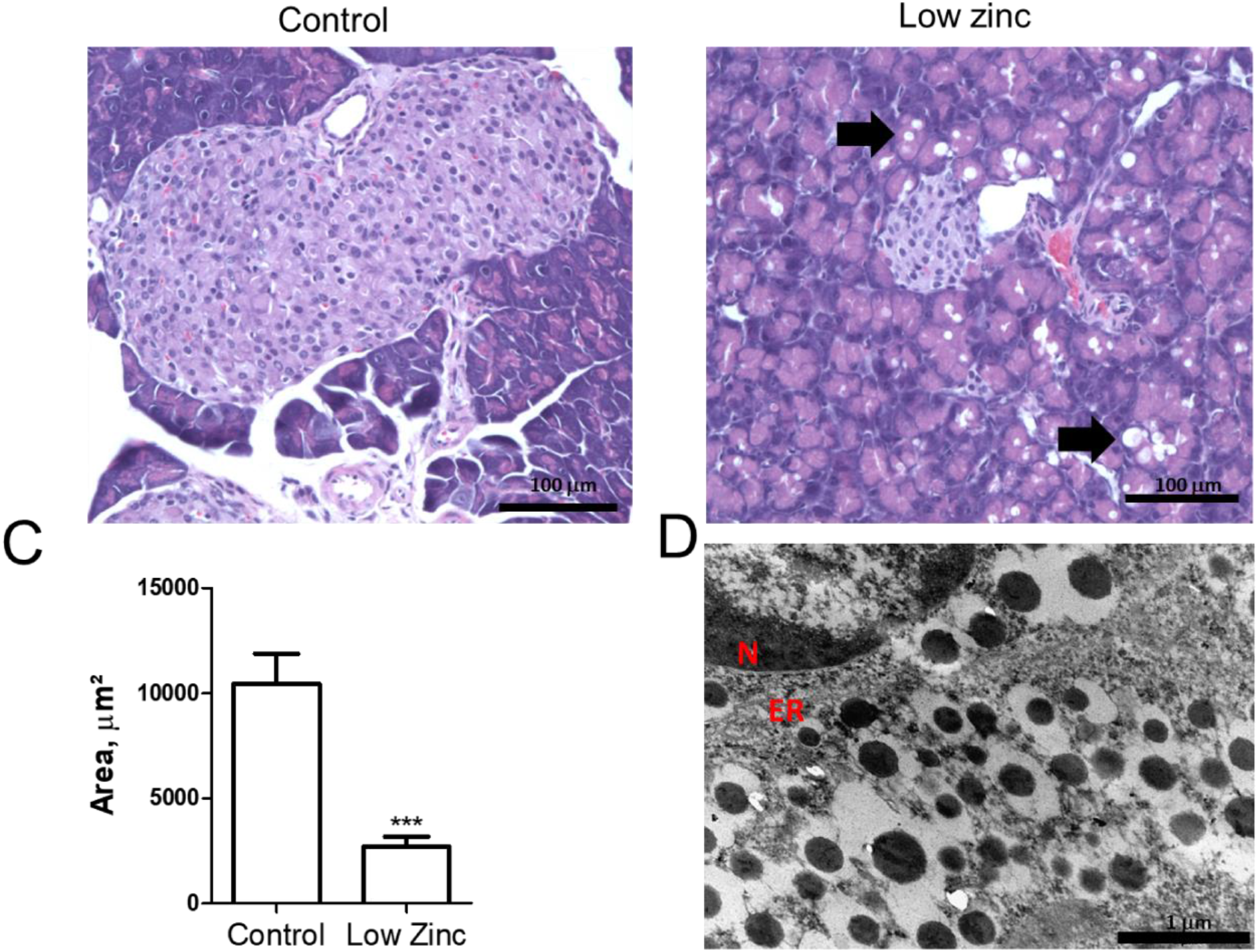
Zinc moderate diet induces pancreatic steatosis and reduction on islet of Langerhans area. Differences between control (A) and zinc deficient (B) diet groups, restrictive diet induces the appearance of adipose tissue on pancreas (black arrow). Islet of Langerhans shows approximately one third smaller on intervention group (C). N = 4 (70 pictures, each group). *** *p*< 0.0001.

Zinc is a well-known cofactor in insulin crystallization [55–57] also within β-cell secretory granules, and mice knock-out for zinc transporter 8 (ZnT8-/-) show no detectable zinc within β-cell accompanied with loss of insulin crystallinity [30]. However, in our present dietary zinc restriction intervention, we found regular insulin crystallinity within pancreatic β-cells, as inferred from ultrastructural data obtained by transmission electron microscopy of the intervention group (**Fig. 2D**), suggesting that body zinc distribution prioritizes insulin crystallinity over other functions.

### Zinc restriction induces islet apoptosis and accumulation of oligomers

As the intervention group showed a reduction in islet size with similar insulinemia and reduced amylinemia, we investigated the β-pancreatic hormonal production. Immunohistochemical analysis of histological cuts revealed a higher labelling for both insulin and amylin in the intervention compared to control group (**Fig. 3**). While we suppose that high insulin production would be compensatory for the higher insulin resistance, the higher pancreatic amylin labeling not followed by higher circulating amylin suggested extra-pancreatic events such as modified metabolization and/or a possible local agglomeration of this hormone. Major amylin accumulation in the islets in the form of amyloid material was discarded giving the lack of evidences by the amyloid stains congo red or Thioflavin S (data not shown).

**Fig 3.**
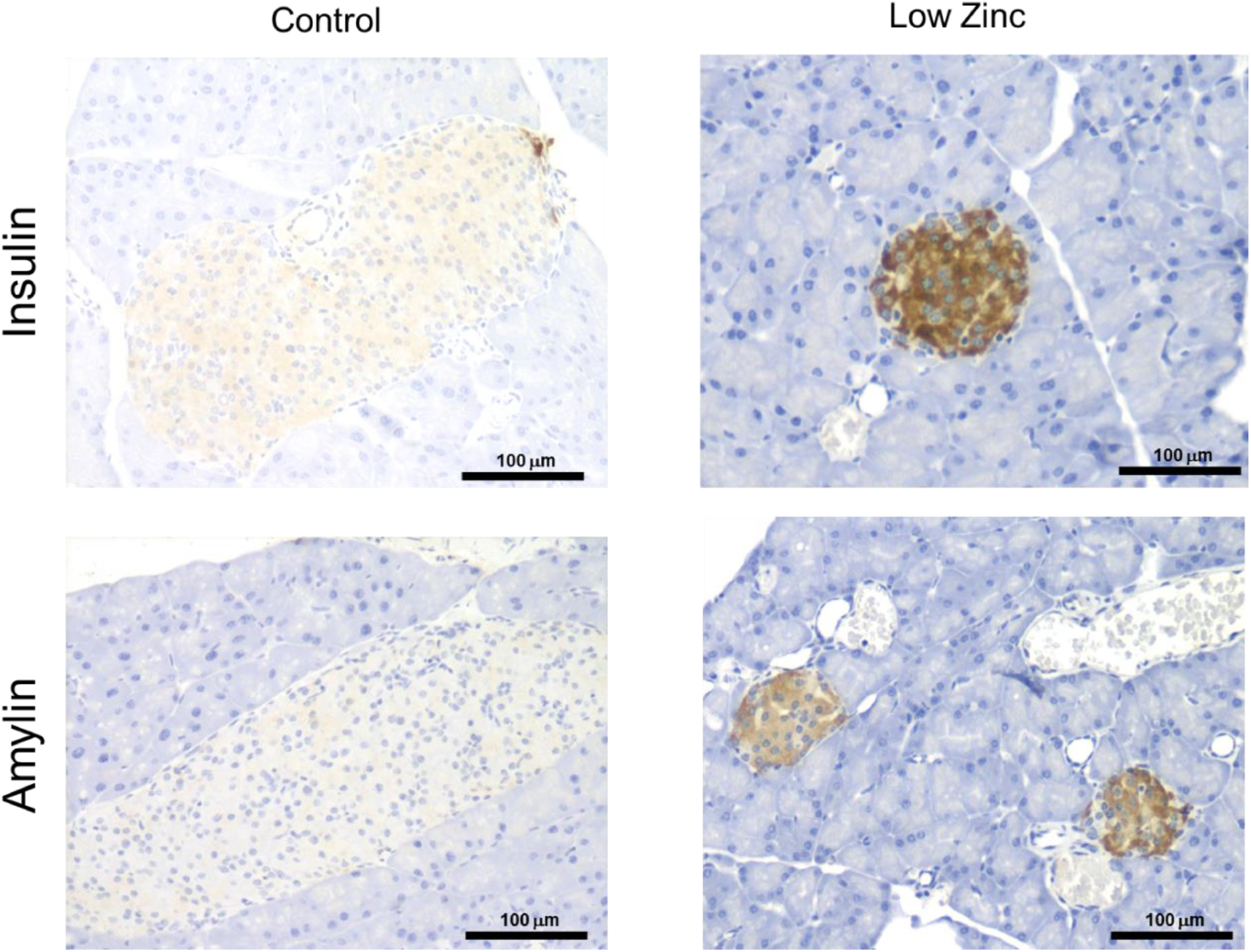
Pancreatic hiperinsulinemia and hiperamilinemia. Immunostaining for insulin (left panel) and amylin (right panel) reveals high local hormonal concentration on intervention group.

In order to test for the presence of oligomers, pancreatic tissues were immunostained for oligomer-like immunoresponsive material (OLIM, A11) [58] (**Fig. 4**), which is selective for oligomeric murine amylin and not responsive to non-agglomerated amylin (**Fig. S11**), in good agreement with previous results showing responsiveness of the OLIM to toxic oligomeric murine amylin material in non-transgenic mouse pancreas [41]. We have detected expressive responsiveness of the OLIM in zinc-restriction dietary intervention group (**Fig. 4B**), while the control diet group showed no labelling (**Fig. 4A**). Successful labeling with A11 was only possible using PBS buffer, while Tris buffer hampered the detection (**Fig. S11**).

**Fig 4.**
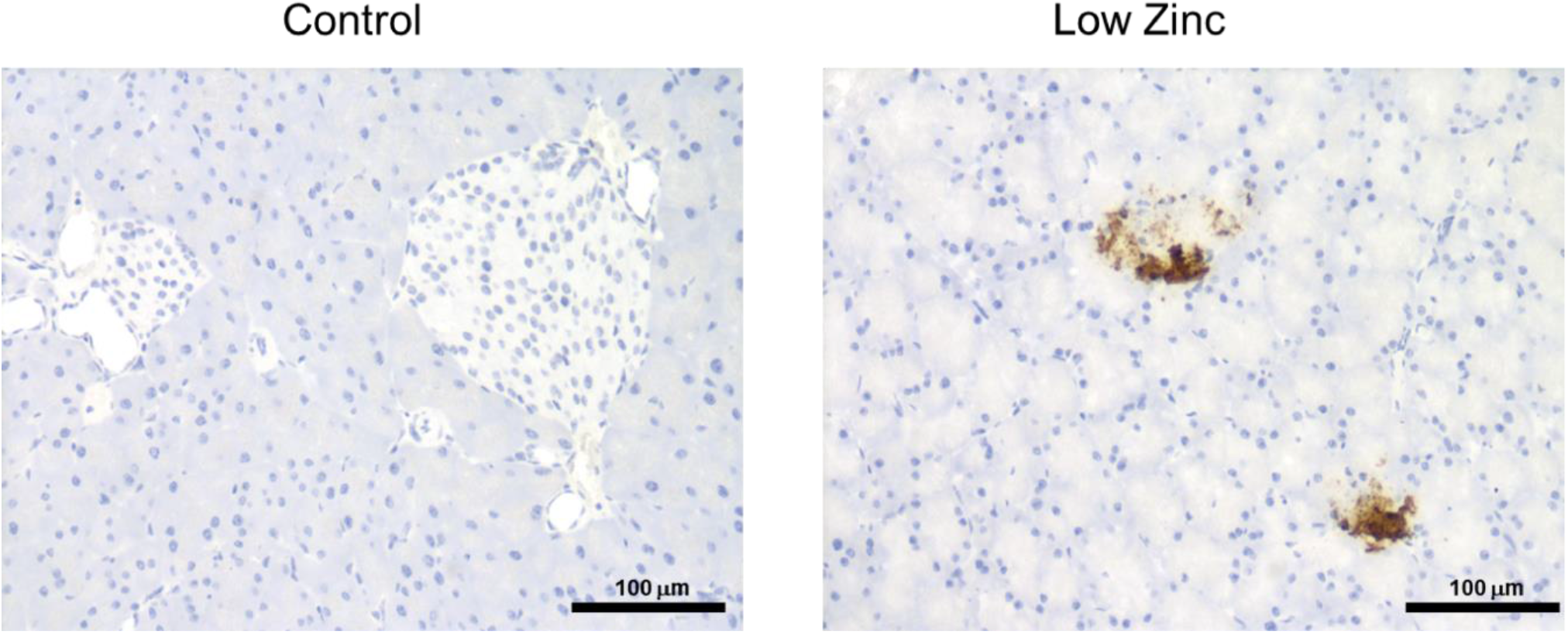
Amylin oligomers on pancreatic tissue in non-transgenic mice. Amylin and oligomers colocalization on pancreatic tissue on hypozincemic group.

Aiming the investigation of the basis for reduced islets size, we tested the pancreas for apoptosis. A TUNEL assay was performed on both groups, revealing positive apoptotic cells in the zinc-restriction group, mostly restricted to the islets (**Fig. 5**), suggesting that zinc-deficient promotes a apoptosis-based mechanism of islet destruction, without apparent relation with pancreatitis, which is characterized by fibrotic lesions and/or thickening vessels. Indeed, total collagen tissue was found reduced on zinc-restricted mice, most likely due to the overall impaired development (**Fig S12)**.

**Figure 5.**
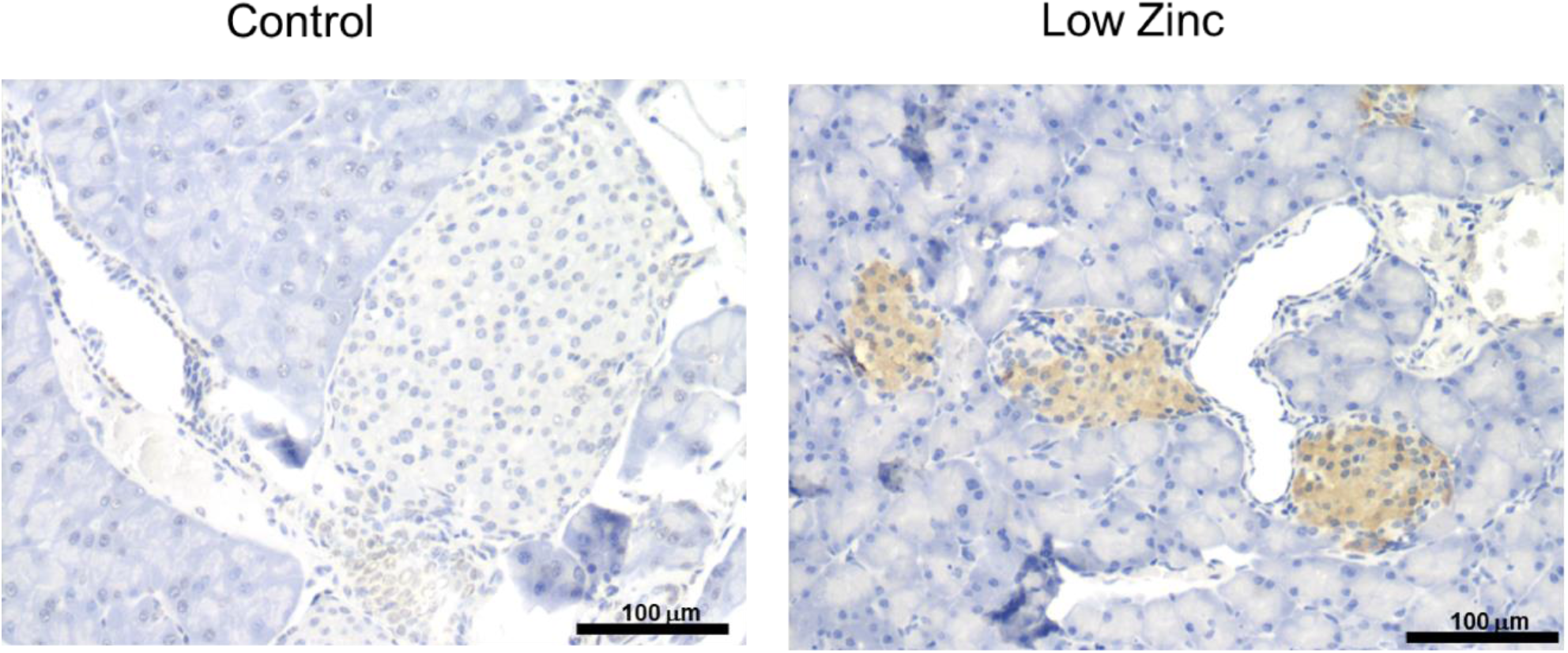
Zinc imbalance induces islet apoptosis. TUNEL staining in control and hypozincemic diet animals.

## DISCUSSION

We have investigated the effect of dietary zinc restriction in the metabolic and pancreatic outcomes in mice. While insulin was still found crystalline within intracellular secretory granules, zinc-restricted animals were clearly impacted by this intervention, resulting in smaller, apoptotic pancreatic islets with high labeling for oligomer-like immunoresponsive material. These data indicate a tight causal effect of zinc restriction in the endocrine pancreas degeneration.

Zinc is an important component of the pancreatic function [59] and it is more concentrated in endocrine cells, mainly in the secretory vesicles of β-cells where stabilizes the structure of insulin granules [55–57,60]. After a stimulus for insulin release, amylin and zinc may be immediately co-distributed locally, but to date no evidence for interaction within the β- granule has emerged. In fact, zinc plays a role in glycemic control [61] and our results suggests that dietary zinc restriction induces a relative insulin resistance (**Fig. 1E**) and impaired endocrine pancreatic function associated with preferable apoptotic pattern despite the insulin crystallinity reached under moderate dietary zinc restriction (**Fig. 1** and **Fig. 3**). Such condition may resemble a subclinical diabetes condition [62,63], characterized by normoglycemic hyperinsulinemia.

Increased insulin levels and insulin resistance are associated with ectopic fat [64,65], and the toxic effect of pancreatic fat accumulation might require a long time before manifesting an impaired β-cell function [66,67]. Pancreatic steatosis was evident in the intervention group (**Fig. 2B**) and islets disorders (**Fig. 3** and **Fig. 4**) may be accelerated due to growth phases with high energy expenditure under nutritional micronutrient deprivation. The decreased plasma amylin concentration (**Fig. 1G**) associated with more prevalent amylin labelling in the islets (**Fig. 3**) along with the presence of oligomers (**Fig. 4**) suggest that modulation on zinc homeostasis impact directly on amylin metabolism and stability, unfolding into toxicity to cells and apoptosis.

For years the study of pancreatic amyloid and oligomers in mice has been hampered by the lack of detection, which supported for long time the concept of non-toxic nature of murine amylin. These limitation stimulated the use of transgenic rodents [68–70]. We have previously reported the *in vitro* amyloid nature of murine amylin [38,39], and subsequently the toxic nature of murine amylin oligomers and its presence in pancreatic tissue in natural abundance of wild-type mice [41]. The techniques used along the years for identification of oligomeric and fibers forms has been demonstrated to have certain limitations [39,71]. In this work, we report that at least part of the limitation in detection by of the OLIM A11 antibody labelling seems to be due to interference of Tris buffer (**Fig. 4** and **Fig. S3**). Tissues washed with Tris buffer displayed similar responsiveness to control or negative control (***Fig. S11***), evidencing the impact of the wash buffer in the responsiveness of the A11 antibody.

Additional studies would be required to provide deeper insight into the mechanisms underlying these processes and perhaps finding biomarkers with clinical relevance. Nonetheless, here, for the first time, we demonstrated that the present experimental model of dietary zinc restriction is enough to independently result in an overproduction of amylin and insulin. These hormones compensate to higher insulin resistance and reduced islet size, along with the formation of a toxic oligomer environment and islet apoptosis in mice.

Although the present data are in view of the thrifty phenotype [72] and the triage [73] theories where micronutrient imbalance seems to exert a key role in the onset of diabetes and comorbidities, the interpretation of this work should not be directly extended to humans or any other *in vivo* system. Instead, it might be used as a hypothesis-generating for such organisms. Another limitation of this work relies in the interpretation for *in vivo* free-living conditions and on the association of other interventions and environmental variables, which might – or not - counteract or enhance the effects rising from zinc-restriction alone.

Further studies are required to determine which cells are under apoptosis, whether autoantibodies are being generated, quantification of other hormones in plasma and pancreas (e.g. MALDI-Imaging) and metals distribution in the organs (by synchrotron X-ray fluorescence). A zinc depletion-repletion approach is also desirable to address the issue on reversibility/management of the pathologic process, addressing whether:

- essential trace mineral micronutrient are independent determinants of a pancreatic (β-cells and others) disfunction/destruction
- zinc and/or other essential trace mineral micronutrients are able to participate in the reversibility/management of pancreatic (β-cells and others) decline through supplementation
- such pancreatic (β-cells and others) decline are followed by elicitation of auto-antibodies and
- tracking early biomarkers that could be proposed for use in diagnosing subclinical process.

The understanding of the relationship between essential trace mineral micronutrient deficiency and the decline in pancreatic β-cells mass could establish the grounds for further human clinical trials, both investigating the biomarkers and mineral status associated to diabetes and the design of reconciliatory therapeutic approaches - nutritional and/or pharmacologic - aimed to at least in part contribute for delay, halt or remission of the decline in pancreatic β-cells mass.

## CONCLUSION

A moderate dietary zinc restriction ensues a metabolism impairment and endocrine disruption with endocrine pancreatic degeneration through apoptosis, involving toxic oligomer formation.

## ACKNOWLEDGMENTS

We would like to thank the Centro Nacional de Bioimagem – UFRJ, the Instituto de Ciências Biomédicas (ICB-UFRJ), Prof. Christina M. Takiya (IBCCF-UFRJ), Profa. Marcia Attias and the Lab for Cellular Ultrastructure Herta Meyer (IBCCF-UFRJ) and their staff for helpful support and providing access to their microscopy facilities, to MSc Daiane Oliveira for helpful assistance with animal experimentation, to the University Hospital Clementino Fraga Filho (HUCFF-UFRJ) for access to the clinical biochemistry platform, and to EMBRAPA for the mineral analysis of the diets.

## DATA AVAILABILITY

The datasets generated during and/or analyzed during the current study are available from the corresponding author on reasonable request.

## AUTHOR CONTRIBUTIONS (according to ICMJE)

Conceived and designed experiments: TS, CKL, LM-A, LMTRL

Performed experiments: TS, CKL, DYS, TMB, NLA, MJA, LM-A, LMTRL

Analyzed the data: TS, CKL, DCS, TMB, NLA, MJA, LM-A, LMTRL

Contributed reagents/materials/analysis tools: TS, CKL, LM-A, LMTRL

Wrote the original draft: TS, LM-A, LMTRL

Revised the manuscript: TS, CKL, DCS, TMB, NLA, MJA, LM-A, LMTRL

Final approval of the version to be published: TS, CKL, DCS, TMB, NLA, MJA, LM-A, LMTRL

