## Supplemental Material for "Zinc restriction promotes β-cell hyper-hormonemia and endocrine pancreas degeneration in mice"

### 1) Composition of control diet – Original Leaflet

**RHOSTER**
INDÚSTRIA E COMÉRCIO LTDA.

MINISTÉRIO DA AGRICULTURA BRASIL ESTABELECIMENTO REGISTRADO SP -08444 ALIMENTAÇÃO ANIMAL

**DIETA ALBUMINA CONTROLE** formulada para roedores em experimentação

| PRODUTOS | % |
|---|---|
| Amido de milho | 39,75 |
| Albumina | 20,00 |
| Amido dextrinizado | 13,20 |
| Sacarose | 10,00 |
| Óleo de soja | 7,00 |
| Celulose MC-101 | 5,00 |
| Mix Mineral AIN-93G | 3,50 |
| Mix Vitamínico AIN-93 | 1,00 |
| L-Cisteína | 0,30 |
| Bitartarato de Colina | 0,25 |
| Tert-Butilhidroquinona | 0,0014 |
| Total | 100,00 |

**AIN-93 Purified Diets for Laboratory Rodents:**
Final Report of the American Institute of Nutrition Ad Hoc
Writing Committee on the Reformulation of the AIN-76A Rodent Diet

**PHILIP G. REEVES, FORREST H. NIELSEN AND GEORGE C. FAHEY, JR.***
United States Department of Agriculture, Agricultural Research Service, Grand Forks Human Nutrition Research Center, Grand Forks, HD 58202-9034 and *Department of Animal Sciences, University of Illinois, Urbana, IL 61801
J. Nutr. 123: 1939-1951, 1993.

**RHOSTER**
INDÚSTRIA E COMÉRCIO LTDA.

MINISTÉRIO DA AGRICULTURA BRASIL ESTABELECIMENTO REGISTRADO SP -08444 ALIMENTAÇÃO ANIMAL

**DIETA ALBUMINA (AIN-93G) RH195115**

| Perfil Centesimal | % | Perfil Calórico | kcal% |
|---|---|---|---|
| Umidade | 9,82 | Proteína | 21,20 |
| Proteína Bruta | 20,08 | Carboidrato | 59,35 |
| Extrato Etéreo | 8,19 | Gordura | 19,45 |
| Fibra Bruta | 1,28 | Total | 100 |
| Cinza | 4,40 | 3,79kcal/g | |
| Carboidrato | 56,23 | | |

**Produtos**

Amido de Milho, Albumina, Amido Dextrinizado, Óleo de Soja, Celulose, L-Cisteína, Mix Mineral AIN-93G, Mix Vitamínico AIN-93, Sacarose, Bitartarato de Colina, Tert-Butilhidroquinona Sigma-Aldrich.

### 2) Composition of Zinc-Restricted Diet – Original Leaflet

RHOSTER
INDÚSTRIA E COMÉRCIO LTDA.

MINISTÉRIO DA AGRICULTURA BRASIL ESTABELECIMENTO REGISTRADO SP -08444 ALIMENTAÇÃO ANIMAL

**DIETA ALBUMINA deficiente em ZINCO** formulada para roedores em experimentação

| PRODUTOS | % |
|---|---|
| Amido de milho | 39,75 |
| Albumina | 20,00 |
| Amido dextrinizado | 13,20 |
| Sacarose | 10,00 |
| Óleo de soja | 7,00 |
| Celulose MC-101 | 5,00 |
| Mix Mineral AIN-93G deficiente em Zinco | 3,50 |
| Mix Vitamínico AIN-93 | 1,00 |
| L-Cisteína | 0,30 |
| Bitartarato de Colina | 0,25 |
| Tert-Butilhidroquinona | 0,0014 |
| Total | 100,00 |

**AIN-93 Purified Diets for Laboratory Rodents:**
Final Report of the American Institute of Nutrition Ad Hoc
Writing Committee on the Reformulation of the AIN-76A Rodent Diet

RHOSTER
INDÚSTRIA E COMÉRCIO LTDA.

MINISTÉRIO DA AGRICULTURA BRASIL ESTABELECIMENTO REGISTRADO SP -08444 ALIMENTAÇÃO ANIMAL

**DIETA ALBUMINA deficiente em Zinco (AIN-93G)** **RH195116**

| Perfil Centesimal | % | Perfil Calórico | kcal% |
|---|---|---|---|
| Umidade | 9,57 | Proteína | 21,10 |
| Proteína Bruta | 20,13 | Carboidrato | 60,22 |
| Extrato Etéreo | 7,92 | Gordura | 18,68 |
| Fibra Bruta | 1,06 | Total | 100 |
| Cinza | 3,86 | 3,82kcal/g | |
| Carboidrato | 57,46 | | |

**Produtos**

Amido de Milho, Albumina, Amido Dextrinizado, Óleo de Soja, Celulose, L-Cisteína, Mix Mineral AIN-93G def Zinco, Mix Vitamínico AIN-93, Sacarose, Bitartarato de Colina, Tert-Butilhidroquinona Sigma-Aldrich.

### 3) Compilation of Diet Composition

| Item | Control | Zinc-restricted |
|---|---|---|
| Corn starch | 39.75 g | 39.75 g |
| Hen egg white solids (dehydrated) | 20 g | 20 g |
| Dextrinized Starch | 13.2 g | 13.2 g |
| Sucrose | 10.0 g | 10.0 g |
| Soy oil | 7.0 g | 7.0 g |
| Cellulose | 5.0 g | 5.0 g |
| Mineral Mix AIN-93 | 3.5 g (*control mix*) | 3.5 g (*Zn-def mix*) |
| Vitamin Mix AIN-93 | 1.0 g | 1.0 g |
| L-Cys | 0.30 g | 0.30 g |
| Choline Bitartrate | 0.25 g | 0.25 g |
| Tert-Butyl Hydroquinone | 1.4 mg | 1.4 mg |

4) Nutritional Data for Powder Pasteurized Hen Egg White Solids (“Albumin”) Used in Present Rodent Diets

http://www.saltosalimentos.com.br/produto_albumina.html

http://www.saltosalimentos.com.br/tabela/Tabela_Clara_Desidratada.pdf

Salto's ALIMENTOS LTDA.

### TABELA NUTRICIONAL
### CLARA DE OVO PASTEURIZADA DESIDRATADA

#### VALOR NUTRICIONAL EM 100g DE CLARA DE OVO

**NUTRIENTES**

| | | |
|---|---|---|
| PROTEINAS | (g) | 84,6 |
| UMIDADE | (g) | 6,5 |
| LIPÍDIOS | (g) | 0,4 |
| CINZAS | (g) | 3,6 |
| CARBOIDRATOS | (g) | 4,8 |
| VALOR ENERGÉTICO | (Cal) | 361,0 |
| COLESTEROL | (mg) | 20 |
| GORDURAS TRANS | (g) | 0 |

**MINERAIS**

| | | |
|---|---|---|
| CÁLCIO | (mg) | 104 |
| FERRO | (mg) | 0,23 |
| MAGNÉSIO | (mg) | 82 |
| FÓSFORO | (mg) | 104 |
| POTÁSSIO | (mg) | 884 |
| SÓDIO | (mg) | 1014 |
| ZINCO | (mg) | 0,135 |
| COBRE | (mg) | 0,128 |
| MANGANÊS | (mg) | <0,03 |

**LIPÍDIOS**

| | | |
|---|---|---|
| SATURADA | (g) | 0 |
| MONOINSATURADA | (g) | 0 |
| POLINSATURADA | (g) | 0 |

**VITAMINAS**

| | | |
|---|---|---|
| NIACINA | (mg) | 0,77 |
| RIBOFLAVINA | (mg) | 3,71 |
| B12 | (mcg) | 0,18 |
| ÁCIDO PANTOTÊNICO | (mg) | 0,67 |
| VITAMINA A | (UI) | <100 |
| TIAMINA | (mg) | <0,05 |
| PIRIDOXINA B6 | (mg) | 0,04 |
| ÁCIDO FÓLICO | (mg) | 0,022 |
| VITAMINA E | (UI) | <0,500 |
| VITAMINA D | (UI) | <90 |

**AMINOÁCIDOS**

| | | |
|---|---|---|
| ALANINA | (mg) | 5160 |
| ARGININA | (mg) | 4920 |
| ÁCIDO ASPÁRTICO | (mg) | 9200 |
| CISTINA | (mg) | 2227 |
| ÁCIDO GLUTÂMICO | (mg) | 11733 |
| GLICINA | (mg) | 3067 |
| HISTIDINA | (mg) | 2063 |
| ISOLEUCINA | (mg) | 4440 |
| LEUCINA | (mg) | 7407 |
| LISINA | (mg) | 5940 |
| METIONINA | (mg) | 3013 |
| FENILALANINA | (mg) | 5163 |
| PROLINA | (mg) | 3260 |
| SERINA | (mg) | 6200 |
| TREONINA | (mg) | 3653 |
| TRIPTOFANO | (mg) | 1437 |
| TIROSINA | (mg) | 3437 |
| VALINA | (mg) | 5763 |

### 5) Quantification of Zinc in Control Diet

LABTEC

**LABTEC LABORATÓRIO DE ANÁLISES QUÍMICAS LTDA**
Rua Equador, 300 - Jd. Santa Clara do Lago II
Hortolândia / SP CEP: 13186.470
www.labtecanalises.com.br

**RELATÓRIO DE ANÁLISE N° 1097205-1**

Cliente: RHOSTER INDUSTRIA E COMERCIO LTDA — Código do Cliente: 1649/24096
Endereço: AV MINISTRO ANTONINO VIEIRA DO AMARAL,1556,JD SALETE
Cidade: ARACOIABA DA SERRA - SP — CEP: 18190-000
CNPJ: 00.984.204/0001-04 — IE: 178.070.220.114
Contato: TERESA BARRETTO

Amostra: DIETA AIN-93G ALBUMINA L:4666/15 - FABR:17/03/2015 — Número da Amostra: 1097205

Data de Recebimento da Amostra: 30/04/2015
Data de Realização do ensaio: 11/05/2015
Data de Emissão do relatório: 14/05/2015

| Análises / Ensaios | Métodos | Resultados | Incertezas | Especificações |
|---|---|---|---|---|
| Zinco (mg/Kg) | ICP OES | 38,6756c | ---------- | --------------------- |

Laboratório: (c) Indica resultado duplamente confirmado.

Maria Cláudia C. Santos
Analista de Laboratório
CRQ - IV Região: 04230901

Maria Helena Bertoni
Responsável Técnico
CRF-SP 12952

Os Resultados referem-se exclusivamente à amostra recebida pelo Laboratório.
Este relatório somente poderá ser reproduzido na íntegra, sua reprodução parcial requer aprovação formal deste laboratório.

Controle Digital: <44144858> **Página: 1/1**

### 6) Quantification of Zinc in Zinc-restricted Diet

LABTEC

**LABTEC LABORATÓRIO DE ANÁLISES QUÍMICAS LTDA**
Rua Equador, 300 - Jd. Santa Clara do Lago II
Hortolândia / SP CEP: 13186.470
www.labtecanalises.com.br

**RELATÓRIO DE ANÁLISE N° 1097204-1**

Cliente: RHOSTER INDUSTRIA E COMERCIO LTDA — Código do Cliente: 1649/24096
Endereço: AV MINISTRO ANTONINO VIEIRA DO AMARAL,1556,JD SALETE
Cidade: ARACOIABA DA SERRA - SP — CEP: 18190-000
CNPJ: 00.984.204/0001-04 — IE: 178.070.220.114
Contato: TERESA BARRETTO

Amostra: DIETA ALBUMINA DEF ZINCO L:4665/15 - FABR:17/03/2015 — Número da Amostra: 1097204

Data de Recebimento da Amostra: 30/04/2015 — Data de Realização do ensaio: 11/05/2015 — Data de Emissão do relatório: 14/05/2015

| Análises / Ensaios | Métodos | Resultados | Incertezas | Especificações |
|---|---|---|---|---|
| Zinco (mg/Kg) | ICP OES | 11,4420c | ---------- | --------------------- |

Laboratório: (c) Indica resultado duplamente confirmado.

Maria Cláudia C. Santos
Analista de Laboratório
CRQ - IV Região: 04230901

Maria Helena Bertoni
Responsável Técnico
CRF-SP 12952

Os Resultados referem-se exclusivamente à amostra recebida pelo Laboratório.
Este relatório somente poderá ser reproduzido na íntegra, sua reprodução parcial requer aprovação formal deste laboratório.

Controle Digital: <44117623> **Página: 1/1**

### 7) Additional data

| A1C% | | | | | |
|---|---|---|---|---|---|
| **Control** | **4.4** | **4.6** | **4.7** | **6.1** | **6.8** |
| **Low Zinc** | **4.1** | **4.6** | **< 4.0** | **< 4.0** | **<4.0*** |

**S1. Reduction of glycated hemoglobin on non-transgenic mice submitted a restricted zinc diet.** After 4 weeks of intervention, both groups – control and zinc-restricted - were evaluated for glycated hemoglobin (A1C), and intervention group exhibited lower A1C compared to control group. N = 5, control group. N= 7, intervention group.

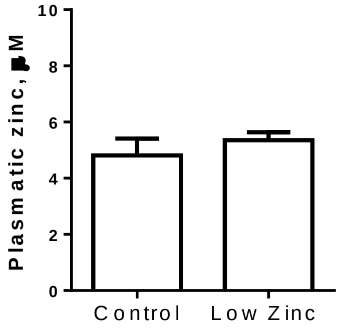

**S2. Zinc plasmatic concentration is not altered by restricted dietary zinc intake.** Weanling Swiss male mice were submitted on a moderate zinc diet for 28 days and no difference on plasmatic zinc content between groups was observed. Zinc was measured by zincon assay. N = 3.

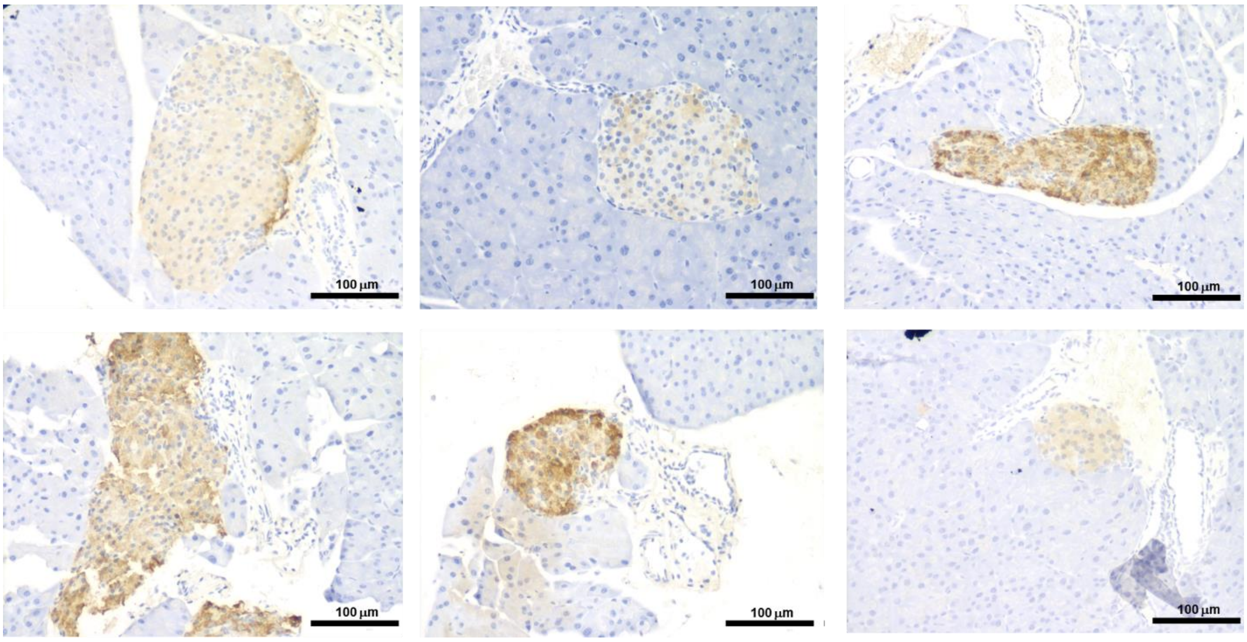

**S3. Pancreatic insulin immunohistochemistry on control group.** Additional representative immunostaining for insulin on control group.

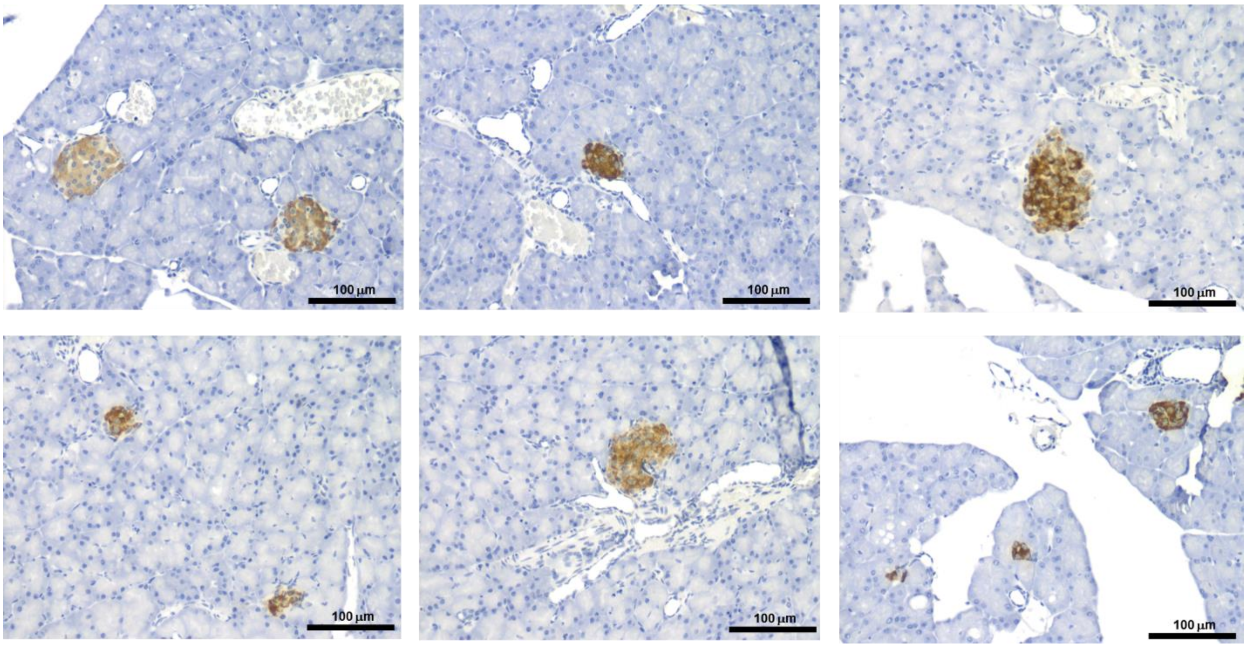

**S4. Pancreatic insulin immunohistochemistry on zinc-restricted diet group.** Additional representative immunostaining for insulin on hypozincemic diet group evidencing intense positive peroxidase staining.

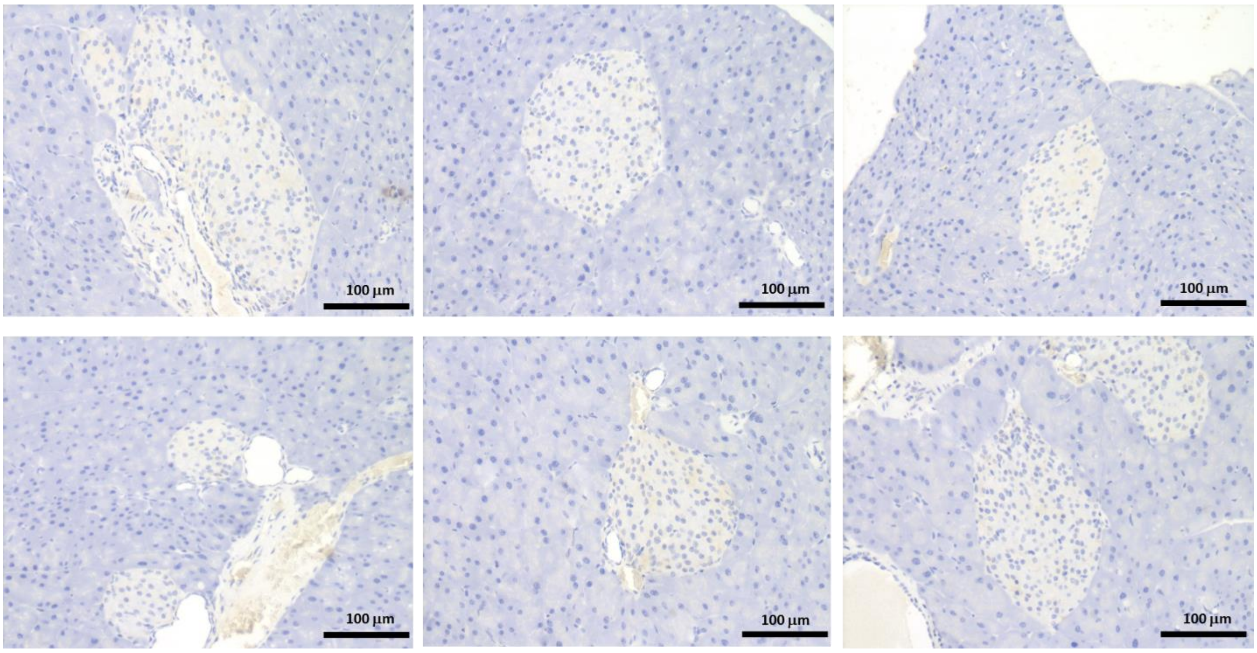

**S5. Pancreatic amylin immunohistochemistry on control group.** Additional representative immunostaining for amylin.

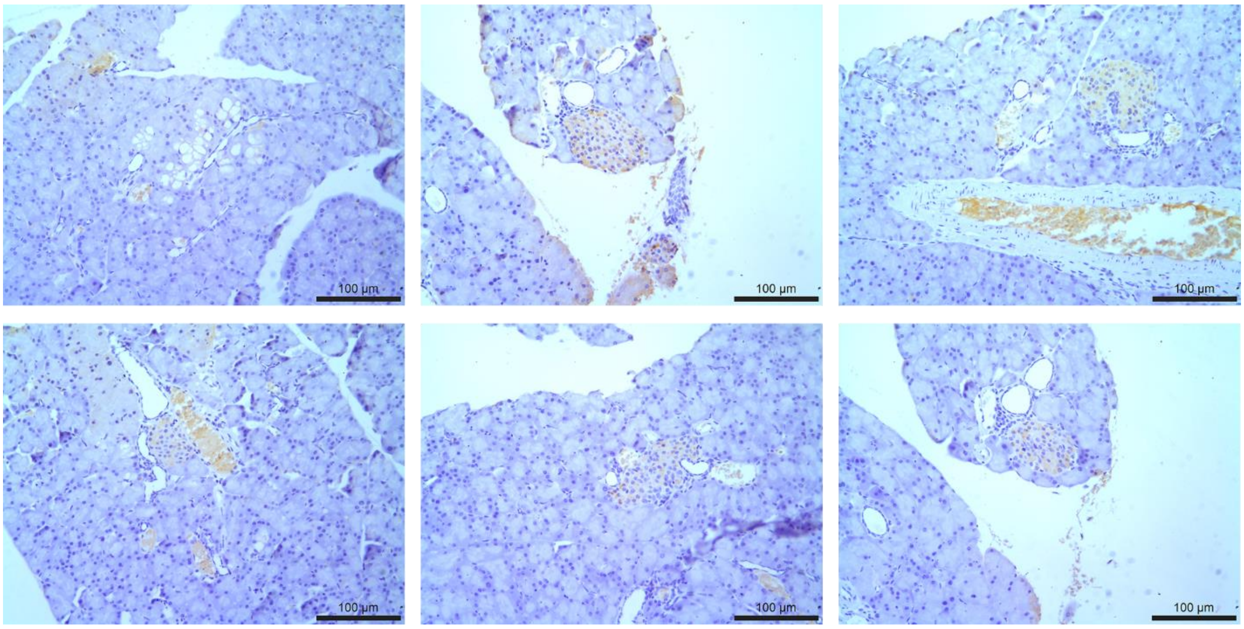

**S6. Pancreatic amylin immunohistochemistry on zinc-restricted group.** Additional representative immunostaining for amylin.

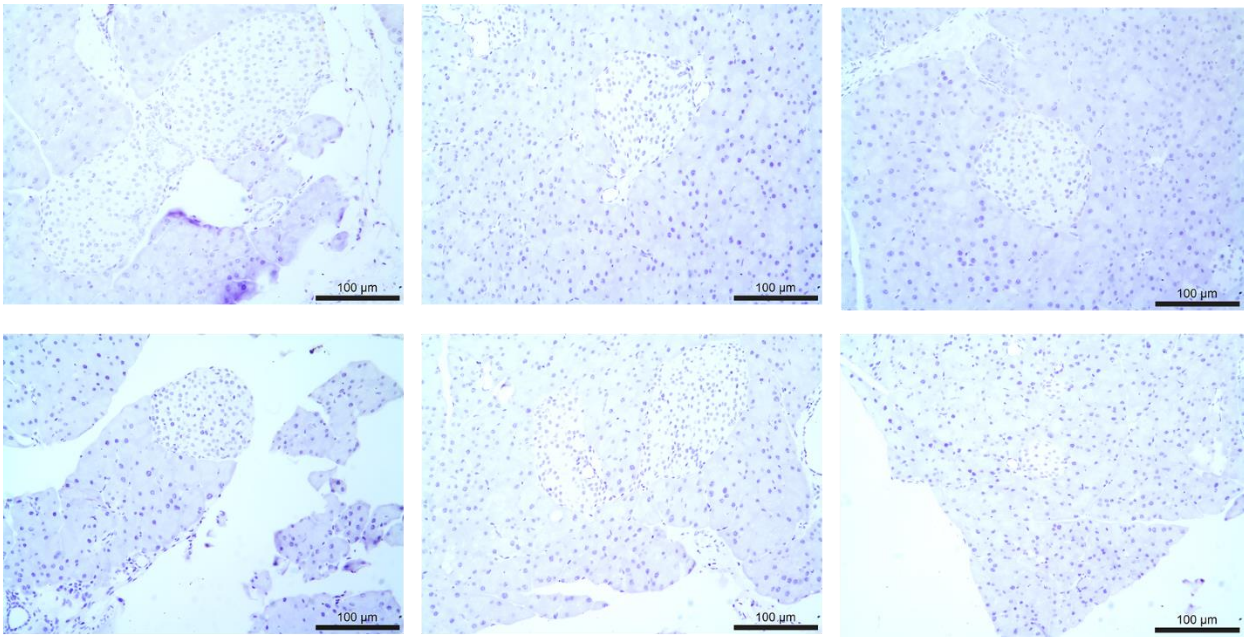

**S7. Lack of detection of responsive material to A11 antibody on islets of control group.** Immunostaining for toxic oligomers species on pancreatic tissue of control group.

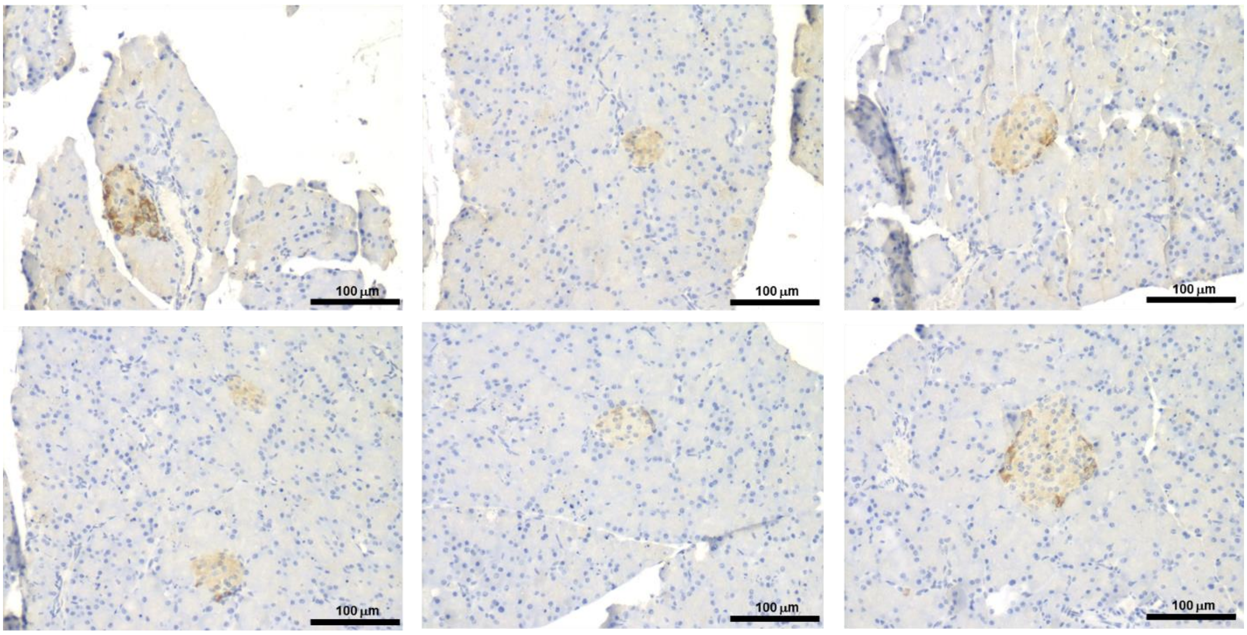

**S8. Positive staining to A11 antibody on islets of zinc-restricted group.** Immunostaining for toxic oligomers species on pancreatic islet of intervention group.

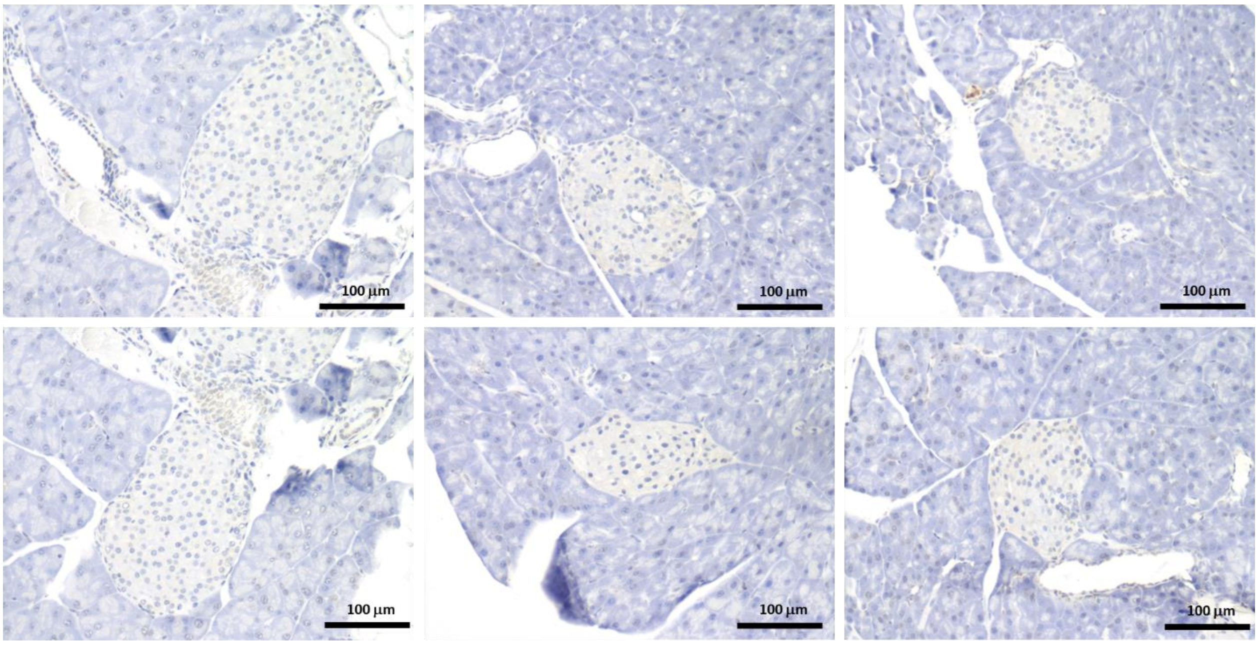

**S9. Additional data for apoptosis assay in islets of control group.** TUNEL assay on control group demonstrating integrity of endocrine and exocrine pancreas.

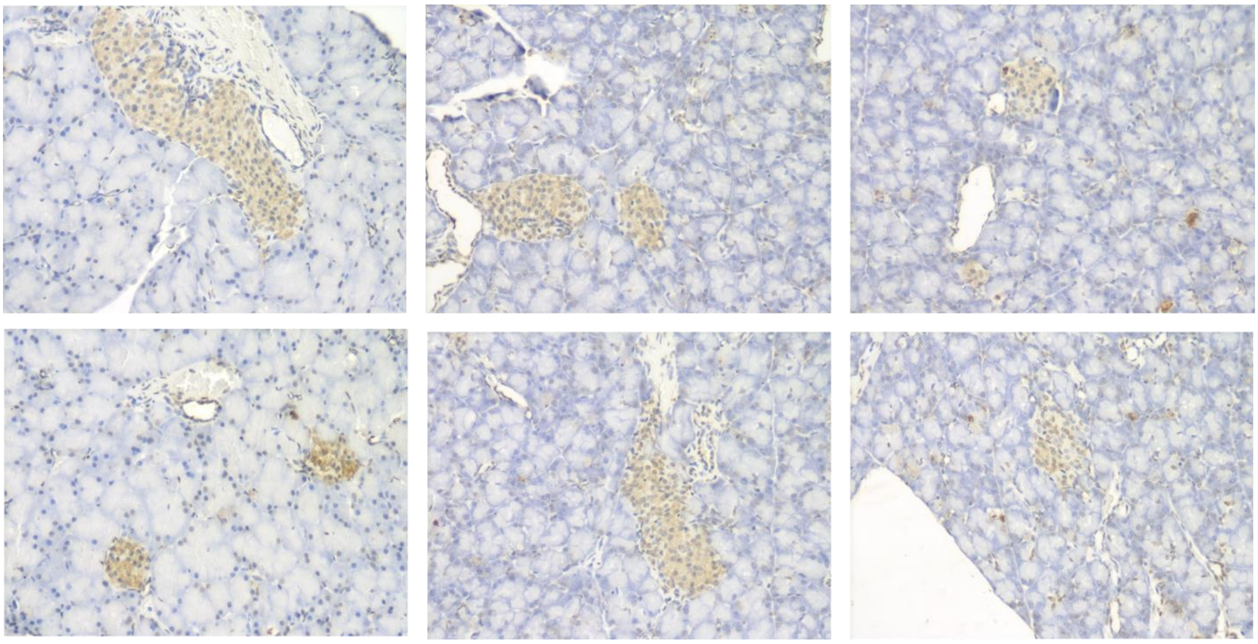

**S10. Additional data for apoptosis assay in islets of zinc-restricted group.** TUNEL assay on intervention group demonstrating genomic instability of endocrine and exocrine pancreas.

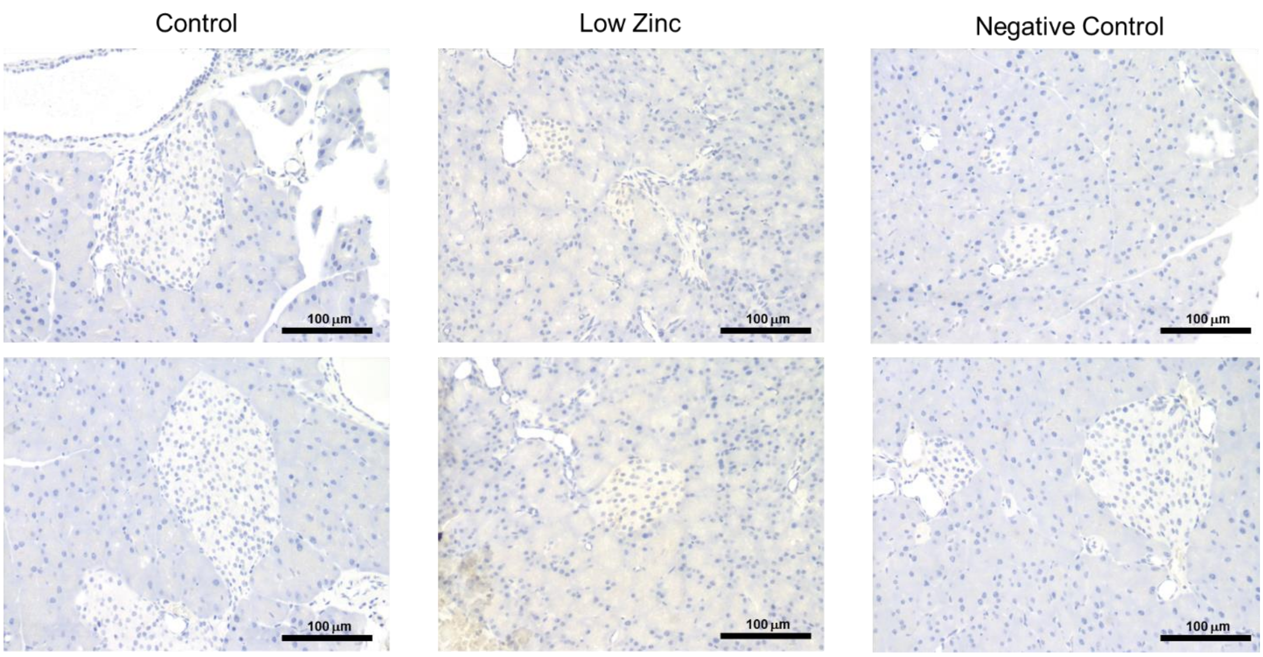

**S11. Effect of buffer in immunostaining for oligomers in islets.** Pancreatic tissue of both groups was washed with 100 mM Tris pH 7.4. Negative control, suppression of the A11 antibody on intervention group, washed with PBS pH 7.4.

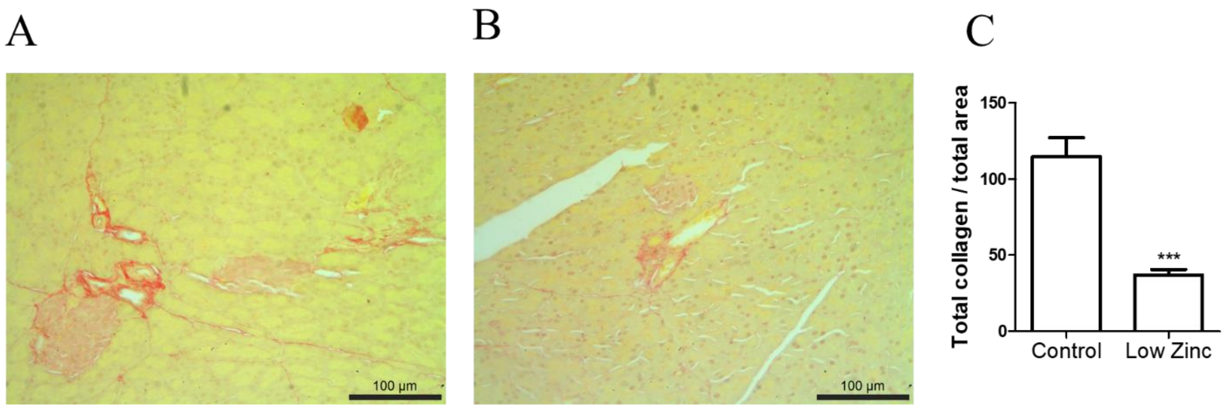

**S12. Collagen quantification on pancreatic tissue.** Sirius red staining on (A) control and (B) intervention pancreatic tissue, quantification of total collagen/total area of tissue on image was performed on Image Pro Plus program (Media Cybernetics Inc. version 6.00.260). N = 70 images/group, ***p<0.0001.
